# Identification of a P62-TIF-IA axis that drives nucleolar fusion and the senescence associated secretory phenotype

**DOI:** 10.1101/2023.12.05.570133

**Authors:** Hazel C Thoms, Tyler Brant, Katie Duckett, Yizheng Yang, Jinxi Dong, Hongfei Wang, Freya Derby, Tumi Akeke, Derek Mann, Fraser R Millar, Alex Von Kriegsheim, Juan Carlos Acosta, Fiona Oakley, Lesley A Stark

## Abstract

Two key characteristics of senescent cells are nucleolar fusion and secretion of a plethora of pro-inflammatory cytokines called the senescence-associated secretory phenotype (SASP). The SASP is dependent on NF-κB but the initial trigger, and links with nucleoli, are unclear. Using multiple *in vitro* and *in vivo* models, we show that an early response to oncogene- and therapy-induced senescence (OIS and TIS) is nuclear/nucleolar accumulation of the PolI complex component, TIF-IA. This accumulation is essential for nucleolar fusion, the SASP and senescence, independent of rDNA transcription. We show that in steady state, TIF-IA is targeted for autophagic degradation by the p62 cargo receptor and that accumulation in senescence occurs as a consequence of ATM activation, which disrupts the p62-TIF-IA interaction. In mice, TIF-IA accumulates in colonic mucosa with age, which is further enhanced in the *nfkb1-/-* model of accelerated ageing. Together, these results reveal a p62-TIF-IA nucleolar stress axis that regulates the SASP and senescence, and that warrants further investigation as an anti-ageing target.

## Introduction

Senescence is complex process that is induced by a wide array of stressors including DNA damage, oncogenes and telomere shortening(1,2). It is a natural process that is essential for development, maintenance of tissue homeostasis, wound healing and cancer prevention (3–5). However, it is a double edge sword as the accumulation of senescent cells in ageing tissue is deleterious to tissue homeostasis, accelerates the ageing process and can promote chronic, age-related diseases including, atherosclerosis, diabetes and cancer (1,6). In addition to a robust cell cycle arrest, senescence is characterised by secretion of an inflammatory milieu known as the senescence associated secretory phenotype (SASP)(7,8). The SASP is a complex and dynamic blend of factors that are key for the initiation and maintenance of cell cycle arrest, paracrine senescence and immunosurveillance (5). The chronic inflammatory environment established by the SASP is considered one of the main mechanisms by which senescent cells impact negatively on tissues to promote ageing and disease(5,9). A master driver of the SASP is the NF-κB transcription factor(7,10,11). NF-κB accumulates on chromatin during senescence and depletion of component proteins can reverse the SASP, without altering cell cycle arrest (10). Although a number of mechanisms have been suggested to explain continued NF-κB activation in senescence (12,13), and cGAS/Sting identified as an important upstream pathway (14,15), the initial triggers of NF-κB signalling upon senescence induction, remain unclear.

Another striking characteristic of senescent cells is gross morphological changes to nucleoli(16–18). The nucleolus is a highly dynamic, membrane-less, nuclear organelle(19,20). In addition to its role in ribosome biogenesis, it acts as a critical stress sensor, coordinating downstream responses to stress such as altered metabolism, cell cycle arrest, DNA repair and apoptosis (21,22). While pre-senescent cells generally have multiple small nucleoli, cells triggered to undergo senescence have one single large nucleolus, which is thought to arise through fusion of phase separated nucleolar droplets (16,18,23–27). Enlarged nucleolar size is also a hallmark of chronological aging and is observed in patients with the ageing disorder, Hutchinson-Gilford progeria syndrome (HGPS) (28,29). While several studies have demonstrated links between these nucleolar changes, ribosome biogenesis and the cell cycle inhibitory effects of senescence (mediated through the canonical, MDM2-P53 nucleolar stress response pathway) (24,30–32), the role of nucleoli in the SASP is unknown.

Previous work from this lab identified a novel, NF-κB nucleolar stress response pathway(22,33). We demonstrated that upon exposure to multiple stress types, the PolI complex component, TIF-IA (RRN3), is degraded in a proteasome and lysosome-dependent manner. Using multiple chemical and genetic modulators, we demonstrated that this degradation is causally linked to a reduction in nucleolar number, an increase in nucleolar area and activation of the NF-κB pathway. Given the similarity between this phenotype and that of senescent cells, we considered that the TIF-IA-NF-kB nucleolar stress pathway may also play a role in triggering the SASP and senescence.

Here we identify TIF-IA as a key regulator of nucleolar fusion and the SASP in senescence. We show that TIF-IA accumulates as an early response to oncogene and therapy induced senescence (OIS and TIS), paralleling nucleolar enlargement and preceding activation of the NF-κB pathway. Inhibiting this accumulation, using siRNA depletion, abrogates nucleolar fusion, the SASP and senescence in OIS and TIS, while overexpression of TIF-IA mimics these effects. We show that normally, TIF-IA is degraded by autophagy in a p62 dependent manner and that during senescence, DNA damage-mediated activation of ATM inhibits the P62-TIF-IA interaction, thereby stabilising TIF-IA. Finally, we demonstrate an association between TIF-IA accumulation and ageing in mice. Together, these data reveal a novel p62-TIF-IA axis that drives nucleolar enlargement, the SASP and senescence. This pathway has significant relevance to ageing and disease.

## Materials and Methods

### Cell lines and treatments

IMR90 human primary fibroblasts stably expressing a 4-hydroxtamoxifen-inducible H-Ras^G12V^-fusion protein (ER:Ras) or a negative control vector (ER:Stop) were a kind gift from Professor JC Acosta (IBBTEC, University of Cantabria), and have been described elsewhere (34). A549 WT and A549-ATG5 ^-/-^ (ΔATG5) cells were a kind gift from Professor Simon Wilkinson (Institute of Genetics & Cancer, University of Edinburgh), whereas WT IMR90 fibroblasts and HCT116 colon cancer cells were obtained from the ATCC. All cells were grown in DMEM (Gibco) supplemented with 10% Foetal Bovine Serum and 1% Penicillin/Streptomycin and were maintained at 37°C in a humidified atmosphere with 5% CO_2_. Treatments were performed under the same conditions. 4-Hydroxytamoxifen (4-OHT) and Etoposide were obtained from Sigma Aldrich, BafilomycinA1 was purchased from Cambridge Bioscience and BrDU was purchased from Abcam.

### Mouse models of aging & senescence

Dr Fraser Millar (Institute of Genetics & Cancer, Edinburgh) kindly provided formalin fixed, paraffin embedded (FFPE) liver sections from mice induced to express either oncogenic Ras (NRas^G12V^), or an inactive mutant (NRas^G12V/D38A^). Full details of how this murine model of oncogene-induced senescence was generated can be found in (35). Briefly, Ras expression (oncogenic or control) was induced in livers of C57BL/6 mice using transposon-mediated intrahepatic gene transfer via a tail vein injection of ‘Sleeping Beauty’ plasmids. Mice were humanely sacrificed 6 days after tail vein injection then tissues collected and processed.

FFPE sections from young (3-4mth) and old (16-17mth) C57BL/6 WT or nfkb1^-/-^ mice were kindly provided by Fiona Oakley (Liver Fibrosis Group, Newcastle University). The nfkb1^-/-^ mice show signs of chronic inflammation and premature aging from 36 weeks of age(36). In all experiments described above, mice were housed in pathogen-free conditions, kept under standard conditions with a 12-hour day/night cycle and free access to food and water. All experiments were approved by the relevant body (Biomedical Research Facility, University of Edinburgh or Newcastle Ethical Review Committee) and performed under a UK Home Office licence in accordance with the ARRIVE guidelines.

### Immunohistochemistry

FFPE colon or liver sections (3μm thickness) were dewaxed and rehydrated using standard procedures. Antigen retrieval was achieved by boiling the slides in 10mM sodium citrate buffer (Fisher), pH 6.0, for 7.5 minutes. Slides were blocked for 30min with 3% H_2_O_2_ (Sigma Aldrich), Avidin/Biotin (Vector Labs) and protein blocking solution (Abcam). They were then incubated overnight at 4°C with primary antibody (Mouse TIF-IA (Abcam), Mouse NRas (SantaCruz), diluted 1:500 in antibody diluent (Dako). Slides were washed with PBS then incubated with biotinylated secondary antibody (1:500) for 30min at room temperature followed by 7.5 minutes of DAB (Abcam). Slides were then dehydrated and fixed with Pertex™ mounting medium (VWR). Immunohistochemistry slides were scanned using the NanoZoomer XR slide scanner (Hamamatsu) with NDP Scan v3.4 software (Hamamatsu). Slides were analysed using QuPath™ Version 0.4.3 software(37). The number of positive cells and the staining intensity was quantified for five equal regions of interest (ROI) per section. Data per ROI were generated and are presented.

### Immunocytochemistry, BrDU assays and image quantification

Cells were grown on glass coverslips then treated as described. Cells were fixed at room temperature for 20min with 4% paraformaldehyde (Thermo Scientific), washed in PBS, then permeabilised for an additional 20min with 0.1% Triton X-100 (Sigma Aldrich). Following fixation, cells were blocked and immunocytochemistry performed using standard procedures (38). Primary antibodies used were TIF-IA (1:200, Assay BioTech B8433), C23 (1:200, Santa Cruz Biotechnology) and p62 (1:100, BD Transductions Labs BD 610833). Alexa Fluor™-conjugated secondary antibodies (Invitrogen) were utilised. Slides were mounted using Vectashield™ medium containing DAPI (Vector Labs). Cell proliferation was determined by analysing the extent of Bromodeoxyuridine (BrDU) incorporation. Briefly, cells were treated with 10μM BrDU for 18 hours prior to fixation and immunocytochemical staining with an anti-BrDU antibody (Invitrogen). Images were visualised with a Zeiss Axio Imager.M2 microscope equipped with an LED light source and captured using an ORCA-Flash 4.0 LT camera (Hamamatsu) with MicroManager™ software to control exposures across the different fluorescent channels. All images per experiment were captured using a constant exposure time. Image analysis was conducted using either FIJI™ (1.54f) or Cell Profiler™ (4.2.5) as specified in figure legends. To quantify nuclear to cytoplasmic ratios of TIF-1A using FIJI™, signal intensity was determined in a defined area of the nucleus (indicated by DAPI) and an equal area at the sub-nuclear periphery (cytoplasm). Nucleolar area was calculated in FIJI™ using the regions devoid of DAPI staining, as previously described(33). Cells exhibiting Senescence-associated Heterochromatic Foci (SAHF) were counted on live images as described elsewhere(39). For all image analysis, at least 100 cells from at least 5 random fields of view were quantified per experiment, for three independent experiments or as specified in the text.

### Plasmids, siRNA and transfections

GFP-tagged WT p62, p62 ΔPB1 and p62 ΔUBA have been described previously (40). pEGFP-C1-TIF-1A WT was kindly gifted by I Grummt (German Cancer Research Centre, Heilderberg). pEGFP-C1-TIF-IA S44A and S44D mutants were generated in house and have been described elsewhere (33). SignalSilence™ SQMTS1/p62 siRNA was obtained from Cell Signalling Technologies. siRNA to TIF-IA was custom made TIF-1A#1 CUAUGUAGAUGGUAAGGUU; TIF-1A#2 CUAGAAUUCCGUUCUUCUA; Control AGGUAGUGUAAUCGCCUUG (33). siRNA was transfected into cells using Lipofectamine 2000 (Invitrogen) according to the manufacturer’s instructions 24h prior to any other treatment. For longer-term knockdown, transfections were repeated as described in the text.

### Quantitative PCR

RNA was extracted from cells using the RNeasy mini kit (Qiagen) following the manufacturer’s instructions. Extracted RNA was purified using RQ1 RNase-free DNase (Promega) then cDNA generated using M-MLV reverse transcriptase and random primers (Promega). Taqman (Thermo Fisher Scientific) or SYBR Green (Roche) assays and a LightCycler 480 system were used to quantify transcript levels. Primers were supplied by Integrated DNA Technologies for the following sequences: *IL-1A For* AGTGCTGCTGAAGGAGATGCCTGA; *IL-1A Rev* CCCCTGCCAAGCACACCCAGTA; *IL-6 For* CCAGGAGCCCAGCTATGAAC; *IL-6 Rev* CCCAGGGAGAAGGCAACTG; *IL-8 For* GAGTGGACCACACTGCGCCA; *IL-8 Rev* TCCACAACCCTCTGCACCCAGT; *MMP1 For*AAAGGGAATAAGTACTGGGC; *MMP1 Rev* CAGTGTTTTCCTCAGAAAGAG; *MMP3 For* TAAAGACAGGCACTTTTGG; *MMP3 Rev* GAGATGGCCAAAATGAAGAG; *NFKB1A For* TCCACTCCATCCTGAAGGCTAC; *NFKB1A Rev*CAAGGACACCAAAAGCTCCACG *TIF-1A For* GTTCGGTTTGGTGGAACTG; *TIF-1A Rev* CGGAATTCTAGCAGCCAGT. The Comparative C_T_ Method (or ΔΔC_T_ Method) was used for calculation of relative gene expression.

### Senescence-associated β-Galactosidase staining

Levels of senescence were determined by examining the extent of β-galactoside activity. Briefly, the cells were fixed with 4% glutaraldehdye in PBS (Sigma Aldrich) for 10 minutes at room temperature. They were then washed with PBS, pH 6.0, and incubated with staining solution (1mg/ml X-Gal (Melford) with 5mM potassium ferrocyanide (Sigma Aldrich) & 5mM potassium ferricyanide (Sigma Aldrich) in PBS) for 16 hours at 37°C in the dark:. Following incubation, the cells were washed with PBS and the percentage of cells staining positive for β-galactoside was determined manually by light microscopy. At least 50 cells were counted from 10 fields of view per replicate.

### Cell lysis and Western blot analysis

Whole cell extracts were prepared by incubating cells on ice with 3 pellet volumes of 1X lysis buffer: Cell signalling 10X whole cell lysis buffer diluted to 1X and supplemented with pepstatinA (Sigma) and PhospSTOP™ (Roche), as per manufacturer’s instructions. Cells were vortexed (3X 30sec) during the incubation period then sonicated (1×10 seconds) using a Soniprep 150 Plus (MSE). Cell debris was pelleted by microcentrifugation at 4°C and protein levels quantified using Bradford Assays (BioRad) as previously described (38). Western blots were carried out using standard protocols as previously described (38). Primary antibodies used were: TIF-IA (rabbit, 1:1000, BioAssayTech B8433); Rrn3 (Mouse, 1:500, Santa Cruz sc-390464); p62 (Mouse, 1:1000 BD Transduction Labs BD610833); GFP (Rabbit, 1:1000 Santa Cruz, sc-8433); Actin-HRP (Mouse 1:2000 sc-47778 Santa Cruz). Bands were quantified using ImageJ as per software instructions.

### Immunoprecipitation assays

Immunoprecipitation assays were performed using 1mg whole cell lysate, prepared in NP40 lysis buffer as previously described (33). Mouse TIF-IA antibody (Santa Cruz Biotechnology) and protein G Dynabeads® (Invitrogen) were to immunoprecipitate endogenous TIF-IA. IgG (pre-immune serum) acted as a control. To immunoprecipitate GFP tagged proteins, cells were transfected with the relevant plasmid, treated as described then whole cell lysates prepared. GFP-Trap® beads (Chromotek)) were used to isolate GFP-tagged proteins. Bound proteins were resolved by SDS polyacrylamide gel electrophoresis then analysed by western blot analysis.

### TIF-IA interactome

Etoposide effects on the TIF-IA interactome were analysed by quantitative mass spectrometry as previously described(41). Briefly, HCT116 cells were treated with DMSO or etoposide (100μM) for 8 hours then pelleted cells disrupted in NP40 buffer and endogenous TIF-IA immunoprecipitated as above. Tryptic peptides were generated by on bead digestion and analysed on a Q-Exactive mass spectrometer connected to an Ultimate Ultra3000 chromatography system (both Thermo Scientific, Germany). Mass spectra were analysed using the MaxQuant Software package in biological triplicate and technical replicate. Fold change for each condition was determined by comparing peptide abundance in GFP V TIF-IA GFP. P values were generated using the data from three replicates.

### Statistical analyses

Graph Pad Prism Software (Version 9) was used for statistical analysis. Normality tests were performed on all datasets to determine the distribution of the data then the relevant parametric or non-parametric test used (as outlined in figure legends) to determine significance. For cell imaging experiments, intensity/size values for all cells analysed in all repeats were pooled, outliers identified and removed using prism software and statistics performed. For analysis of mouse sections, Tukeys multiple comparison test was performed on QuPath data from five ROI per mouse per condition (see above). For quantitative PCR, P values were generated by comparing three technical repeats, for three individual experiments, or as specified in the figure legend. The number of repeats for all experiments are given in the figure legends and the exact P values specified. Data were deemed to be significant if P<0.05.

## Results

### 2.1 TIF-IA accumulates upon senescence induction *in vitro* and *in vivo*

To determine the role of TIF-IA-NF-κB nucleolar stress in senescence, we initially used the well characterised model of OIS in which IMR90 human diploid fibroblasts, transduced with an Oestrogen receptor:H-RAS^G12V^ fusion protein (ER:RAS), are exposed to 4-hydroxytamoxifen (4-OHT) so that they undergo a coordinated OIS response (34)(Fig.S1A). Firstly, we explored TIF-IA levels and found that there was a significant *increase* in TIF-IA protein three days post RAS^G12V^ induction, particularly in the nucleus/nucleolus (Figs. 1A and S1B). This increase occurred as an early response to OIS induction, paralleling nucleolar fusion (as indicated by elongated nucleolar morphology and increased nucleolar area) but, preceding activation of the NF-κB pathway, as determined by transcription of the early NF-κB gene target, NF-κB1A (IκB) (Figs 1B to D and S1B and C).

**Figure 1.**
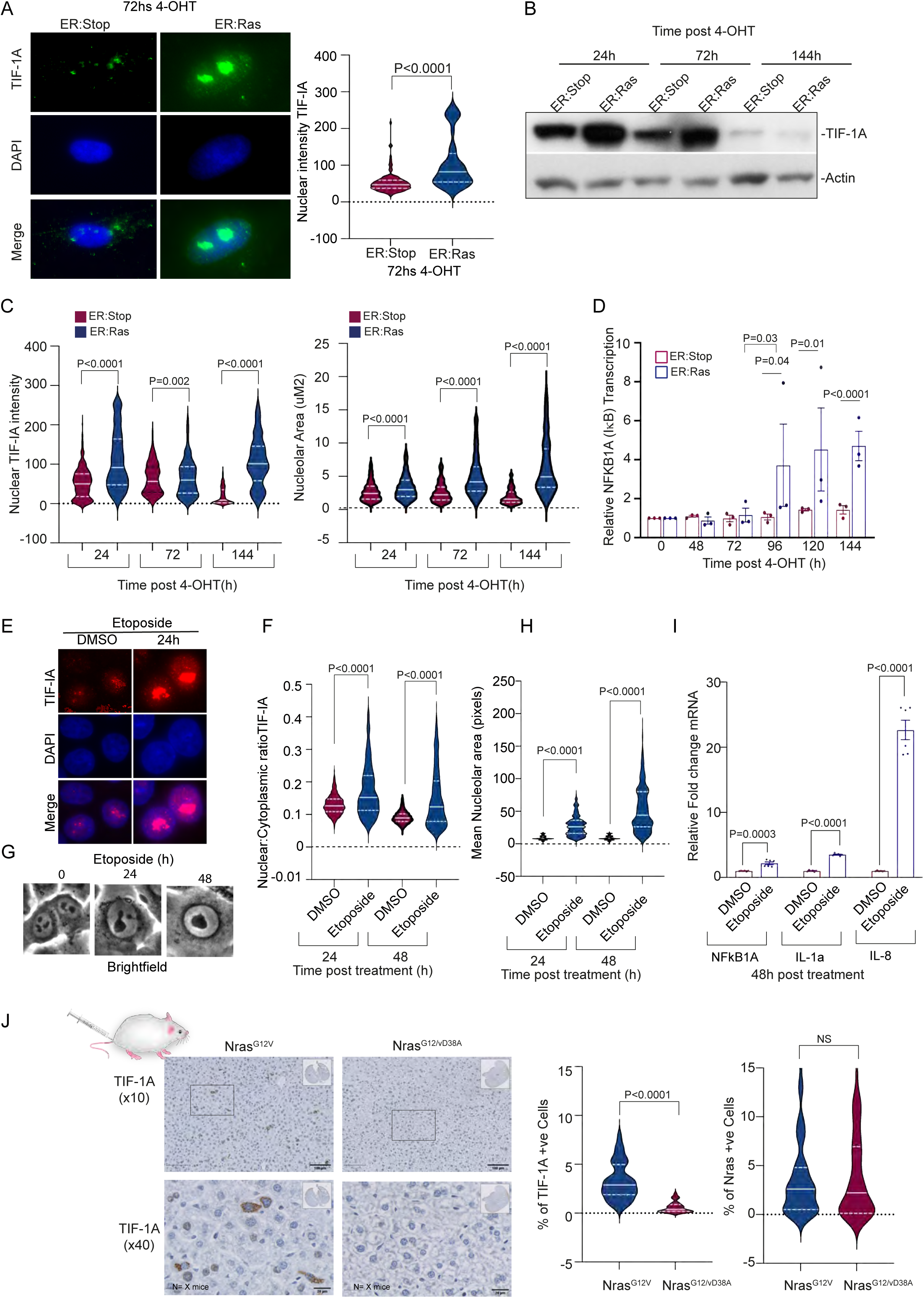
TIF-1A protein accumulation is an early event in oncogene and therapy-induced senescence. (A to D) TIF-IA accumulation parallels nucleolar fusion and precedes NF-κB activation in oncogene-induced senescence (OIS). IMR90 ER:Ras cells were treated with 4-hydroxytamoxifen (4-OHT) for the times specified to induce oncogenic Ras expression. IMR90 ER:Stop cells serve as a control and retain proliferative capacity with 4 -OHT. (A) Immunomicrographs showing nucleolar (areas devoid of DAPI staining) accumulation of TIF-IA in ER:Ras cells 72hs post 4-OHT exposure. (B, C) Levels and localisation of TIF-IA were monitored in ER:Stop and ER:Ras cells exposed to 4-OHT in time course studies using (B) Western blot analysis, performed on whole cell extracts. See Fig S1C for quantification. N=3 (C) Immunocytochemistry. See Fig S1D for representative images of full time course. FIJI™ software was used to quantify nuclear TIF-IA intensity and nucleolar area (areas devoid of DAPI staining). At least five fields, minimum of 150 cells, were analysed per experiment. N=5. (D) qRT-PCR was used to monitor expression of early NF-κB target gene, NFKB1A (IκB), in ER:Stop and ER:Ras cells treated with 4-OHT for the times given. (E to I) Nucleolar accumulation parallels nucleolar fusion and precedes transcription of SASP factors in therapy-induced senescence (TIS). HTC116 cells were treated with etoposide (100uM) for the times specified in hours (h). (E and F) Immunocytochemistry was used to determine TIF-IA levels and localisation. (E) Representative immunomicrographs. (F) Cell Profiler™ was used to quantify the intensity of cytoplasmic and nuclear TIF-1A and the ratio for each cell determined. Five fields of view (minimum 80 cells) per treatment were quantified for each time point (n=3). (G) Representative phase-contrast images of live HCT116 cells treated with 100μM Etoposide for the times indicated (*x40* magnification). Nucleolar fusion is evident. (H) Nucleolar area was determined from phase contrast images using FIJI™ in at least 500 nuclei/treatment. (n=2). (I) Expression of the indicated NF-κB target genes was quantified using qRT-PCR. The bars represent the mean ± the standard error (n=3). (J) TIF-IA and N-Ras immunohistochemistry were performed on liver sections from mice culled 6 days after hydrodynamic delivery of Nras^G12V/D38A^ (*n* =5) or Nras^G12V^ (*n* =5) transposons. TIF-IA accumulation is observed in oncogenic Nras^G12V^ expressing, but not control Nras^G12V/D38A^ expressing, hepatocytes (left violin plot), while N-ras expression is similar (right violin plot). Top panel: representative images at *10X* (scale bar 100uM) and *40X* (scale bar 20uM) magnification. Ten regions of interest (ROIs) were randomly selected from each slide and the intensity of TIF-1A and N-Ras staining quantified using QuPath™ software (v0.2.3). The left violin plot shows the percentage of TIF-1A positive cells for each ROI, whereas the right plot exhibits NRas staining. There were 5 mice analysed from each group. Statistical significance throughout was determined using one-way ANOVA, with Tukey’s multiple comparison, Mann Whitney U test or Student’s t-Test, as appropriate. Ns-non-significant.

To validate these data, we used a model of therapy-induced senescence (TIS) where human IMR90 fibroblasts, or epithelial cancer cells (HCT116), were induced to become senescent by addition of etoposide, a topoisomerase inhibitor known to induce DNA double-strand breaks. Senescence is observed in this model using 48h treatment with etoposide followed by 6 days standard medium (Fig. S1E). Similar to OIS, we found that nuclear accumulation of TIF-IA was an early response to TIS, occurring within the first 24h of treatment (Figs 1E and S1F). Brightfield images clearly demonstrate TIF-IA accumulation is paralleled by nucleolar fusion (elongated structure) at 24h followed by the emergence of a single prominent nucleolus and activation of the NF-kB target genes and SASP factors, IkB (NFkB1A), Il-1a and IL-8 (Figs 1E to I).

Next, we explored the *in vivo* relevance of our findings using a murine model in which OIS is induced in hepatocytes via hydrodynamic delivery of a mutant Nras^G12V^ encoding transposon, along with a sleeping beauty transposase expressing plasmid. A plasmid encoding an Nras^G12V^ effector loop mutant, incapable of downstream Nras signalling (Nras^G12V/D38A^), was used as a negative control. Sections of livers from these mice were provided by the Acosta lab who previously demonstrated that 6 days post transduction, there is a significant increase in senescence markers (*dcr2* and *arf)* and SASP factors (IL-1B) in livers of mice expressing Nras^G12V^, compared to those expressing the control Nras^G12V/D38A^ (35). Similarly, we found that 6 days after transduction there was a significant increase in expression of TIF-IA in livers of mice expressing the oncogenic Nras^G12V^, compared to those expressing the Nras^G12V/D38A^ control (Fig. 1J). Analysis of Nras demonstrated that an equitable percent of hepatocytes expressed Nras^G12V^ and TIF-IA (Fig 1J). It also demonstrated a similar expression of Nras^G12V^ and Nras^G12V/D38A^ (Fig. 1J). Together, these data indicate that TIF-IA accumulates in response to senescence induction *in vitro* and *in vivo,* which is associated with emergence of a single large nucleoli and transcription of NF-kB target genes.

### TIF-IA is required for OIS and TIS

To further explore the link between TIF-IA accumulation, nucleolar size and the SASP, we depleted the protein prior to and during OIS induction (Fig 2A). One of the initial events in OIS is hyperproliferation. It occurs within 24h of oncogene induction and is associated with DNA damage and consequently, the SASP. Given TIF-IA is essential for rDNA transcription, we considered that increased TIF-IA may be required for the hyper-proliferative phase of OIS. However, BrDU incorporation assays indicated that depleting TIF-IA prior to senescence induction had no effect on RAS^G12V^ – induced hyper-proliferation, or indeed, on the subsequent cell cycle arrest (Fig. 2B). Similarly, depletion had no effect on the formation of senescence associated heterochromatin foci (SAHF), which are linked to DNA damage (Fig. 2C). In contrast, immunocytochemistry, using the nucleolar marker C23, indicated TIF-IA depletion significantly blocked the increase in nucleolar area in OIS. Loss of TIF-IA also inhibited transcription of SASP factors, as indicated by qRT-PCR (Figs 2D and E) and the reinforcement of senescence, as indicated by reduced β-galactosidase (SA-β-Gal) activity (Fig. 2F).

**Figure 2.**
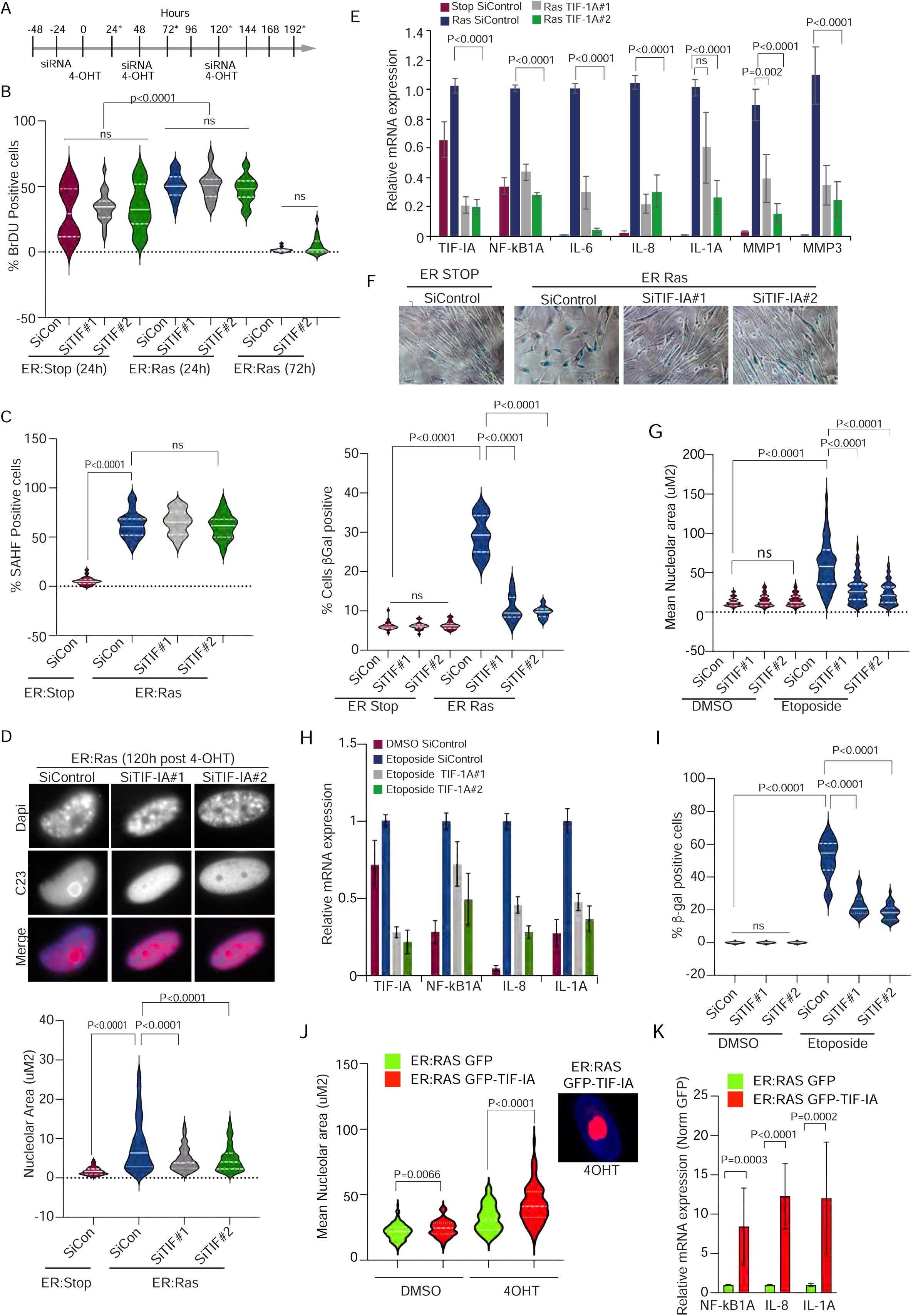
TIF-IA depletion attenuates the SASP and senescence in OIS and TIS. A to F IMR90 ER:Stop or ER:Ras fibroblasts were pre-treated with two independent TIF-IA siRNA (SiTIF-1A#1 & SiTIF-1A#2) or a non-sense sequence (SiControl) prior to addition of 4-OHT for various times. (A) Schematic showing the treatment schedule for siRNA and 4-OHT delivery. (B) BrDU incorporation was used to assess cell proliferation. BrDU positive cells were stained by immunocytochemistry and quantified manually by fluorescent microscopy. At least 100 cells from at least 10 fields of view were analysed for each treatment group (n=2). (C) The percentage of cells with senescence associated heterochromatin foci (SAHF) (large, bright DAPI foci) was determined by microscopic analysis of the DAPI channel 120h after 4-OHT exposure, with a minimum of 200 cells counted manually per treatment for each replicate (n=2). (D) Anti-C23 (nucleolar marker) immunocytochemistry. Top panel: representative images from 4-OHT treated cells. Bottom panel: quantification of nucleolar area (measured as areas devoid of DAPI using FIJI™ software) for at least 50 cells per condition (n=2). (E) qRT-PCR demonstrating TIF-IA depletion inhibits Ras-induced transcription of SASP factors. Induction in ER:Ras siControl cells was compared to that in ER:Stop SiControl cells. Expression in ER:Ras, TIF-IAsiRNA treated cells is given as a percentage of the induction in SiControl cells for each gene and expressed as the mean ± SEM (n=3). (F) IMR90 cells were treated with 4-OHT for 8 days and the effect of TIF-1A knockdown on senescence was determined. Top:Typical fields of view after treatment. Below: the percentage of β-Galactosidase positive cells was quantified in 10 fields of view per replicate by light microscopy (n=3). (G to I) HCT116 cells were transfected with control or two independent TIF-IA siRNAs (#siTIF-IA 1, #siTIF-IA 2) prior to treatment with 100uM Etoposide for 48h (G and H) or 72h (I). Depletion of TIF-1A mRNA was assessed by qRT-PCR (H). (G) Live images were acquired as in Figure 1G and nucleolar area determined using FIJI™ software. At least 125 nucleoli per treatment were assessed from 3 individual replicates. (H) Expression of TIF-IA, IκB (NF-KB1A) and the SASP factors, IL-1α and IL-8 was quantified by qRT-PCR. Bars represent the relative expression compared to etoposide siControl +/-SE (n=3). (G) β-galactosidase assays were performed on etoposide treated cells. The percentage of senescent cells (indicated by blue stain) was determined by light microscopy. Ten fields of view, at least 50 cells/field, were counted for each experimental condition. N=3. (J) IMR90 ER:Ras cells were transfected with pEGFP-C1 or pEGFP-TIF-IA, treated for 5 days with DMSO or 4OHT then immunocytochemistry performed for the nucleolar marker, C23 (inset shows example immunomicrograph). Nucleolar size was quantified using FIJI™ software and area of C23 stain. 33-100 cells were imaged per replicate (n=3). (K) HCT116 cells were transfected with eGFP-C1 or eGFP-TIF-IA. qRT-PCR was used to determine the levels of the given NF-κB target genes 72h later. GFP expression (relative to GAPDH) was used to normalise for transfection efficiency. Statistical significance throughout was determined using either one-way ANOVA, with Tukey’s multiple comparison, Mann Whitney or Student’s t-Test, dependent on normality.

Similar to OIS, siRNA depletion of TIF-IA abrogated the effects of the therapeutic agent etoposide on nucleolar area, transcription of SASP factors and SA-β-Gal activity (Figs 2 G to I). We conclude from these data that in response to senescence induction, TIF-IA accumulates in nuclei/nucleoli downstream of DNA damage, which is causally associated with gross morphological changes to nucleoli and transcription of SASP factors. In support of this conclusion, overexpression of TIF-IA in IMR90 fibroblasts caused an increase in nucleolar area, which was further enhanced by RAS^G12V^ induction (Fig. 2J). It also caused an increase in expression of IκB and SASP factors (Fig. 2K).

### The role of TIF-IA in OIS and TIS is independent of rDNA transcription

The relationship between senescence, ribosome biogenesis and downstream effector pathways is complex and remains controversial. While many studies have demonstrated an increase in rDNA transcription upon OIS and replicative senescence (24,42,43), others have found a decrease(32,44). These alterations in ribosome biogenesis have been linked to p53 stabilisation and cell cycle arrest. However, few studies have explored links to the SASP. Therefore, we next considered whether accumulation of TIF-IA induces the SASP and senescence by modulating rDNA transcription.

qPCR for the 47S pre-RNA transcript demonstrated that the accumulation of TIF-IA post ER:RAS induction, and the subsequent activation of NF-κB, is actually associated with a significant reduction in 47S rDNA transcription, (Figs 1C and D and 3A). Mimicking this reduction, using siRNA to TIF-IA, did not mimic effects of OIS on the SASP or SA-β-Gal activity (Figs 3B and 2F). Similarly, etoposide treatment of IMR90 cells caused TIF-IA accumulation and nucleolar enlargement alongside decreased 47S transcription (Fig 3C and S1F). In contrast, etoposide-mediated TIF-IA accumulation in HCT116 cells was associated with *increased* 47S transcription, paralleled by nucleolar enlargement and transcription of SASP factors (Figs 3C and 1E and F). Together, these data would suggest that the effects of senescence induction on 47S transcription are context dependent. They also suggest that while nucleolar fusion and induction of the SASP in OIS and TIS is dependent on TIF-IA accumulation, there is no link with rDNA transcription. At this point we cannot rule out links to altered rRNA processing.

**Figure 3.**
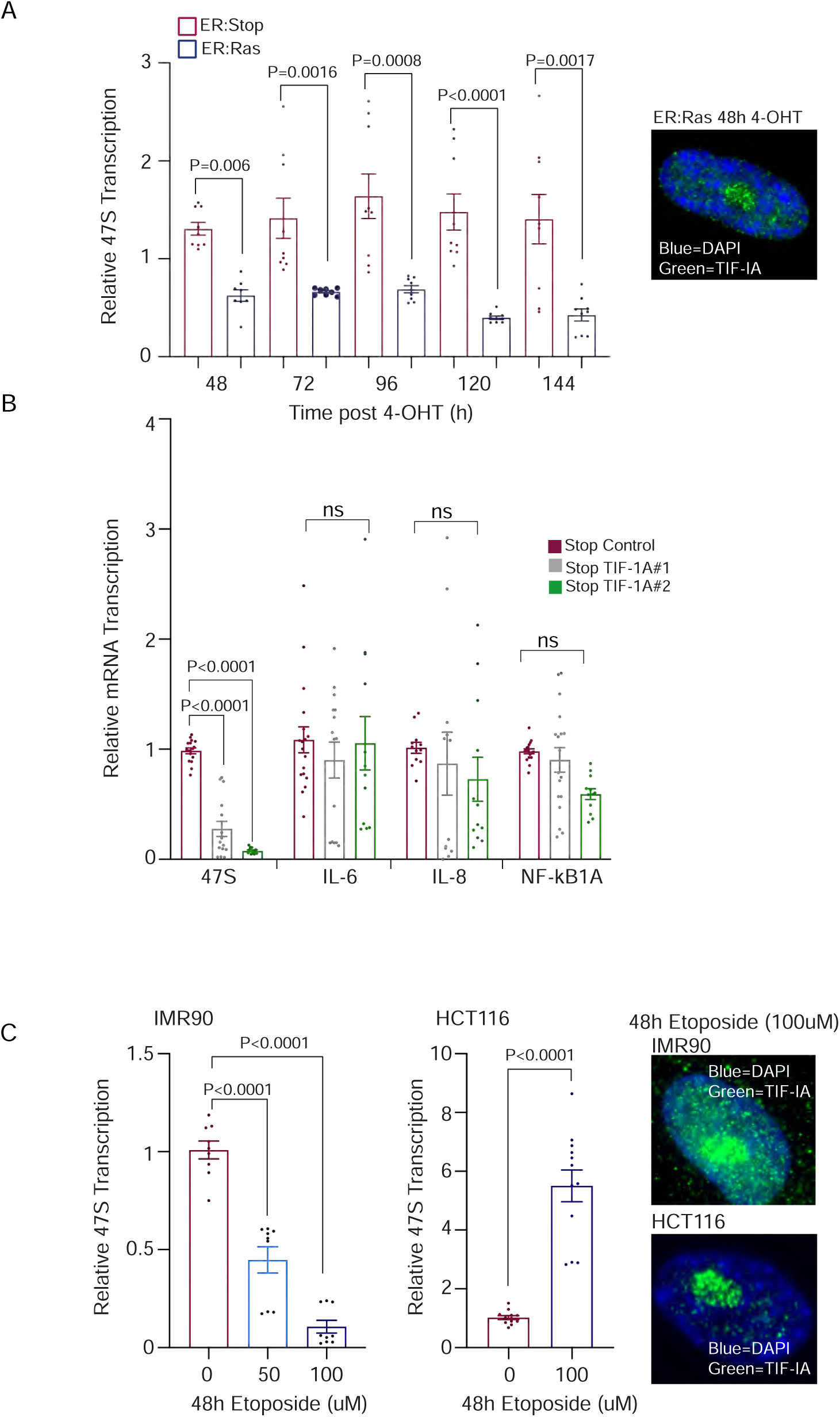
Role of TIF-IA is independent of rDNA transcription. (A) TIF-IA accumulation is associated with reduced 47S transcription in IMR90 cells. qRT-PCR was used to monitor 47S transcription in ER:Stop and ER:Ras cells treated with 4OHT for the times given. Each point represents a technical replicate (n=3) (B) Depleting TIF-IA alone reduced 47S transcription but has no effect on transcription of SASP factors. siRNA was used to deplete TIF-IA in ER:STOP cells as in figure 2 then qRT-PCR used to monitor expression of the given NF-κB target genes 144h later. Each point represents a technical replicate (n=5) (C) qRT-PCR was used to monitor 47S transcription in IMR90 (n=3) and HCT116 (n=4) cells treated with the given concentrations of etoposide for 48h. Right: Representative immunomicrographs showing nucleolar accumulation of TIF-IA. Statistical significance throughout was determined using a students T test.

### P62-mediated autophagic degradation of TIF-IA prevents senescence

OIS and TIS induction had a minimal effect on TIF-IA mRNA (Figs. 4A and 2H), suggesting that the early accumulation in pre-senescence is due to protein stabilisation. Given the importance of this accumulation in nucleolar fusion and the SASP, we set out to understand the mechanisms involved in regulating TIF-IA stability to prevent senescence, and the pathways triggered to induce it.

**Figure 4.**
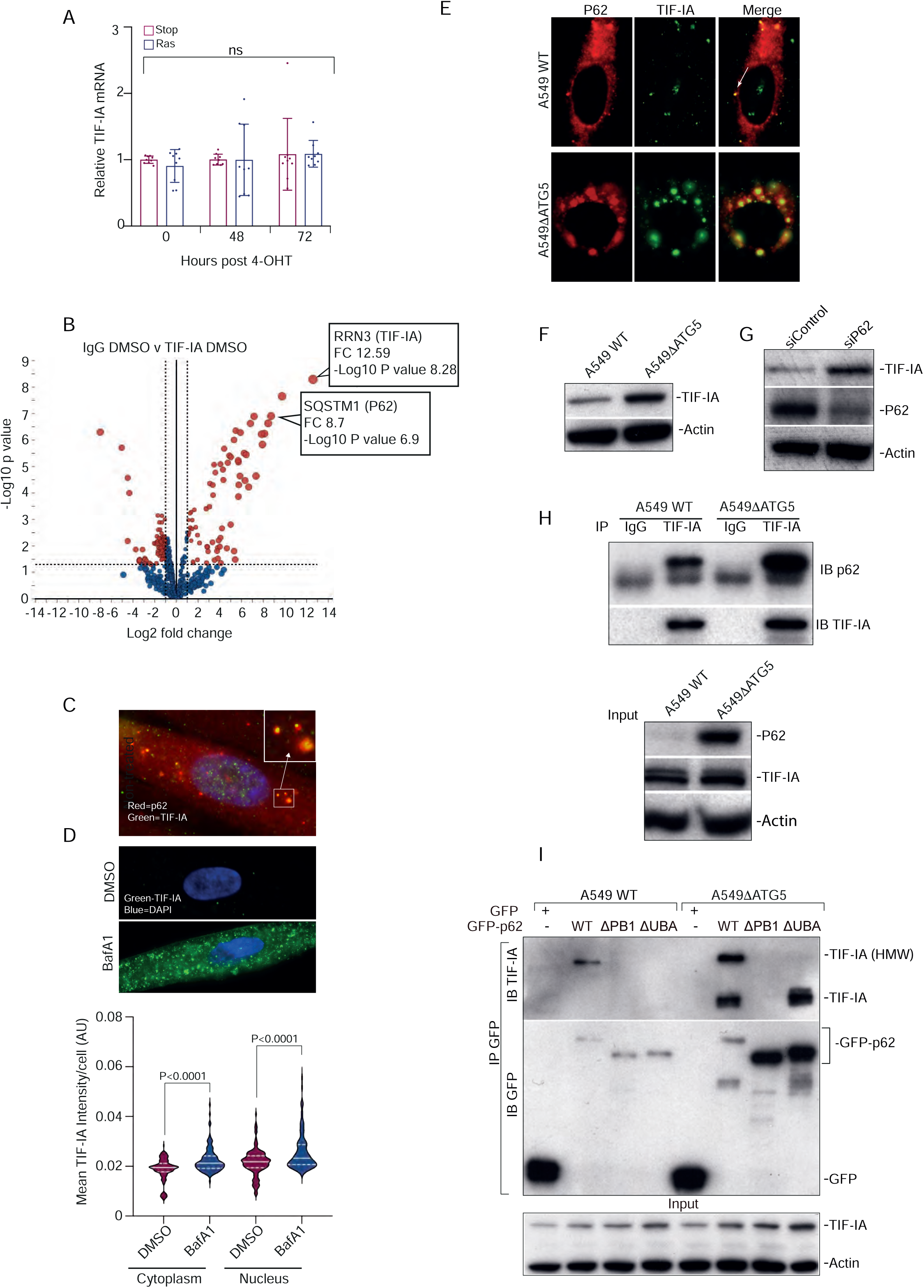
p62 binds to and regulates the stability of TIF-IA. (A) The levels of TIF-1A transcription were investigated by qRT-PCR in IMR90 ER:Stop and ER:Ras cells exposed to 4-OHT for the times indicated in hours (n=3). (B) HCT116 cells were treated with DMSO (control) or etoposide for 8h. Whole cell lysates were immunoprecipitated with IgG or TIF-IA antibody then precipitated proteins analysed by quantitative mass spectrometry. Volcano plot shows a comparison of IgG v TIF-IA for DMSO treated cells. Blue dot=non-significant. Red dot=significant change (fold change (FC) >2, P<0.05). (C) Immmunomicrograph showing p62-TIF-IA co-localisation in cytoplasmic foci in IMR90 cells. (D) Top: Immunomicrograph of IMR90 cells treated with the lysosome inhibitor, bafilomycin A (BafA1) or DMSO carrier. Bottom: Cell profiler was used to quantify cytoplasmic and nuclear TIF-IA intensity. A minimum of 50 cells were analysed/treatment for each biological repeat. N=2. (E) Representative immunomicrographs (63X) showing level and localisation of TIF-IA and p62 in wild type (WT) A549 cells and those in which the autophagy gene, atg5, has been deleted (ΔATG5). White arrow indicates co-localised foci in WT cells. N=5. (F) Immunoblot performed on whole cell extracts from WT and Δatg5 cells. n=5 (G) Immunoblot showing levels of TIF-IA and p62 in whole cell lysates following siRNA depletion of p62. n=2. (H) TIF-IA was immunoprecipitated (IP) from WT or Δatg5 A549 cells then recovered proteins analysed by immunoblot (IB) for p62. Stripped gels were re-probed for TIF-IA. Input levels of protein are shown. Rabbit IgG acts as a control. n=3 (I) P62 binds to a high molecular weight (HMW) form of TIF-IA in a manner dependent on the dimerization and ubiquitin binding domains. Wild type and Δatg5 A549 cells were transfected with plasmids expressing GFP-P62 WT, GFP-p62 ΔPB1 (deleted for the PB1 domain required for dimerization) or GFP-p62ΔUBA (deleted for the ubiquitin binding domain). GFP-tagged proteins were immuoprecipitated (IP) from whole-cell lysates using GFP-TRAP beads. Precipitated proteins were subjected to immunoblotting (IB) for TIF-IA and GFP. Input levels of TIF-IA are shown. N=2. Actin is used as a loading control throughout.

Firstly, pull down assays and quantitative mass spectrometry were employed to identify TIF-IA binding proteins in control cells, and during TIS initiation. HCT116 cells were treated with DMSO (control) or etoposide (100uM. 8h). TIF-IA antibody or IgG control were used to pull TIF-IA protein complexes out of total cell lysates, then complexes analysed by mass spectrometry. The 8h time point was chosen as it was the earliest at which TIF-IA accumulation was evident, and preceded NF-κB pathway activation (Fig S2A and B). Analysis of these mass spectrometry data revealed that one of the top TIF-IA interacting proteins was the autophagy cargo receptor, SQSTM1 (P62) (Fig. 4B). TIF-IA was the top interacting protein, confirming the validity of the methodology (Fig. 4B). The detailed results of the interactome studies are provided in Table S1. The interaction with P62 was of particular interest in terms of TIF-IA stability and so, we investigated this further.

Immunocytochemistry confirmed that TIF-IA co-localised with p62 in cytoplasmic puncti in IMR90 fibroblasts, and that levels of TIF-IA increased in the presence of the lysosomal inhibitor known to block autophagy: bafilomycin A1 (a selective inhibitor of the vacuolar-type H+–adenosine triphosphatase) (Figs 4C and D). Similarly, TIF-IA co-localised with p62 in discrete cytoplasmic foci in wild type A549 cells, which was significantly enhanced by deletion of the autophagy component, ATG5 (Fig 4E). Total cell levels of TIF-IA were elevated in ATG5 deleted cells and in cells in which p62 was depleted using siRNA (Figs 4F and G). Immunoprecipitation experiments confirmed that TIF-IA binds to p62 in control cells, and that the levels of TIF-IA, and the TIF-IA-p62 interaction, are enhanced by autophagy inhibition (Fig. 4H). Further mapping of the p62-TIF-IA interaction revealed wild type p62 binds to both a native and a high molecular weight (HMW) form of TIF-IA (Fig. 4I). Binding to both forms is lost upon deletion of the p62 PB1 domain (which is responsible for homodimerization and PKCa binding) while binding to HMW TIF-IA is lost upon deletion of the ubiquitin binding domain (UBA) (Fig. 4I). Together our data suggest that under basal conditions, TIF-IA is targeted for autophagosomal degradation by p62 in a manner dependent on ubiquitin binding, and that this degradation prevents senescence.

### P62-TIF-IA binding is lost in senescence in a manner dependent on the DNA damage response kinase, ATM

Senescence is generally associated with increased autophagy (50). Given that we observed an increase in TIF-IA protein in OIS and TIS, rather than a decrease, we speculated that the p62-TIF-IA interaction is lost upon senescence induction. Indeed, immunoprecipitation assays demonstrated reduced p62-TIF-IA binding as an early response to TIS (Fig. 5A). As expected, there was a reduction in P62 levels when TIS was triggered, in keeping with enhanced autophagy. However, quantification of IP and input immunoblots demonstrated the reduction in p62-TIF-IA binding was more pronounced than the reduction in input p62 alone, suggesting reduced binding was not merely a consequence of reduced p62 protein (Fig. 5A). Etoposide did not appear to alter P62-TIF-IA binding in the interactome studies, which may be explained by the earlier time point used (Table S1).

**Figure 5.**
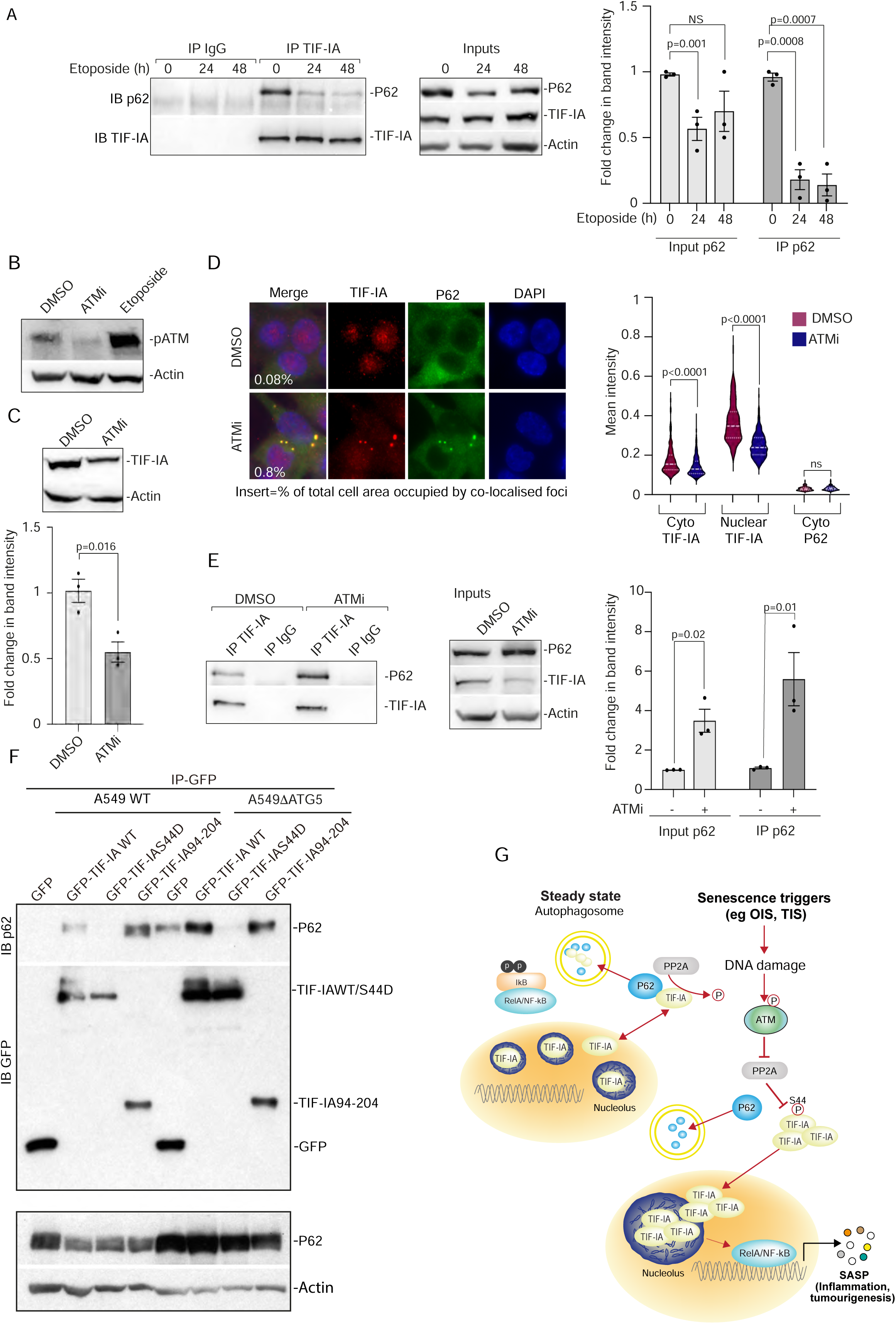
TIF-IA-p62 binding is modulated in senescence downstream of DNA damage. (A) TIF-IA was immunoprecipitated (IP) from HCT116 cells treated with etoposide (100uM) for the indicated times then recovered proteins analysed by immunoblot (IB) for p62. Stripped gels were re-probed for TIF-IA. Input levels of protein are shown. Rabbit IgG acts as a control. ImageJ was used to quantify p62 intensity for input (relative to actin) and IP samples. N=3. (B) Anti-phosphorylated ATM (pATM) immunoblot performed on whole cell lysates from HCT116 cells treated with a small molecule ATM inhibitor (ATMi) (10h), etoposide (24h) or carrier control (DMSO). (C) HCT116 cells were treated with ATMi for 8h then western blot analysis performed on whole cell lysates using the indicated antibodies. Top:representative immunoblot. Bottom: ImageJ was used to quantify TIF-IA band intensity relative to actin. N=3. (D) Left: Immunomicrographs demonstrating level and localisation of TIF-IA in HCT116 cells treated with carrier (DMSO) or ATMi. ImageJ was used to quantify the percentage of cell area occupied by yellow foci. Right: Cell profiler was used to quantify mean intensity of the indicated proteins in the cytoplasm (cyto) and nucleus. N=2. (E) HCT116 cells were treated with DMSO or ATMi for 8h. TIF-IA was immunoprecipitated and precipitates immunoblotted for p62 and TIF-IA. Left: representative immunoblots. Input levels are shown. Rabbit IgG acts as a control. Right: ImageJ was used to quantify band intensities as in A. (C) Model of suggested TIF-IA senescence pathway. In steady state, TIF-IA shuttles between the nucleolus, nucleoplasm and cytoplasm. PP2A dephosphorylates S44 allowing TIF-IA ubiquitination, P62 binding and autophagic degradation. Upon senescence induction, DNA damage activates ATM which inhibits PP2A activity, TIF-IA S44 dephosphorylation and p62 binding. TIF-IA consequently accumulates which causes nucleolar fusion, NF-kB activation and the SASP. See the discussion for further details Actin acts as a loading control throughout. Statistical significance determined by Students T-Test.

Ataxia-telangiectasia mutated kinase (ATM) is a DNA damage response (DDR) kinase that is activated upon induction of TIS and OIS (Fig 5B) (45). Of all the DDR kinases, ATM appears to be particularly important for the SASP and senescence(46). Although ATM activation contributes to increased autophagy in senescence, it can also inhibit autophagic degradation of specific proteins following senescence induction(47). Using the KU55933 ATM inhibitor (ATMi) (Fig. 5B), we explored the role of this kinase in TIF-IA degradation.

Immunoblot and immunocytochemical analysis indicated that ATMi caused a significant reduction in cellular levels of TIF-IA protein (Fig 5C and D). Furthermore, immunoprecipitation assays and IHC demonstrated an increase in TIF-IA-P62 binding and co-localisation of the two proteins in distinct cytoplasmic foci upon ATM inhibition (Figs 5D and E).

Protein phosphatase 2A (PP2A), which dephosphorylates TIF-IA on serine 44, is a direct target of the ATM kinase (48,49). This is of interest as we have previously shown that TIF-IA protein accumulates upon exposure to the PP2A inhibitor, okadaic acid(33). We have also demonstrated that mutating S44 of TIF-IA to the phospho-mimetic, aspartic acid (S44D), completely blocks stress-mediated degradation of the protein (33). Based on these data, we considered that S44 dephosphorylation is required for TIF-IA-p62 binding, which is inhibited downstream of ATM activation. Indeed, immunoprecipitation assays demonstrated that the TIF-IA S44D mutant is unable to bind p62 (Fig. 5F). Further deletion studies revealed that amino acid residues 94-204 of TIF-IA are sufficient for the p62 interaction (Fig. 5F).

Based on the above and previous data, we propose a model whereby under normal conditions, TIF-IA is targeted for autophagic degradation through its association with p62. However, during senescence, ATM-mediated inhibition of PP2A causes an accumulation of TIF-IA phosphorylated at S44. This in turn causes loss of p62 binding, translocation of the protein to the nucleus/nucleolus and consequently, nucleolar fusion, stimulation of the NF-κB pathway and transcription of SASP genes. Defining the mechanism by which TIF-IA accumulation causes nucleolar fusion and NF-κB pathway activation is of incredible interest, but is out with the scope of this manuscript.

### TIF-IA accumulates in colonic tissue during mouse ageing

Loss of the Nfkb1 (p50) gene in mice leads to early onset ageing and a reduction in overall lifespan(36,50). Up to 6 month of age, Nfkb1^-/-^ mice are comparable to their wild type counterparts. However, by 12 months they demonstrate distinctive, age-related phenotypes including increased DNA damage, low-grade inflammation, aggravated cell senescence and impaired regeneration in the liver and gut.

We used young (3-4 months) and older (16-17 months) wild type (WT) and Nfkb1^-/-^ mice to explore TIF-IA in chronological and genetically induced mouse aging. Quantitative immunohistochemistry revealed that for both wild type and Nfkb1^-/-^ mice, there was a significant increase in nuclear TIF-IA in the gut of older mice, compared to the same organ of young mice (Figs 6A and B). Comparing WT to Nfkb1^-/-^ mice, we found no significant difference in gut TIF-IA levels in young mice, in keeping with their similar phenotype at this age. However, there was a highly significant increase in TIF-IA in the gut of older Nfkb1^-/-^ mice, compared to the gut of older WT mice, consistent with the increased DNA damage and inflammation observed in Nfkb1^-/-^ mice (Figs 6A and B). Thus, TIF-IA accumulates during ageing *in vivo*.

**Figure 6.**
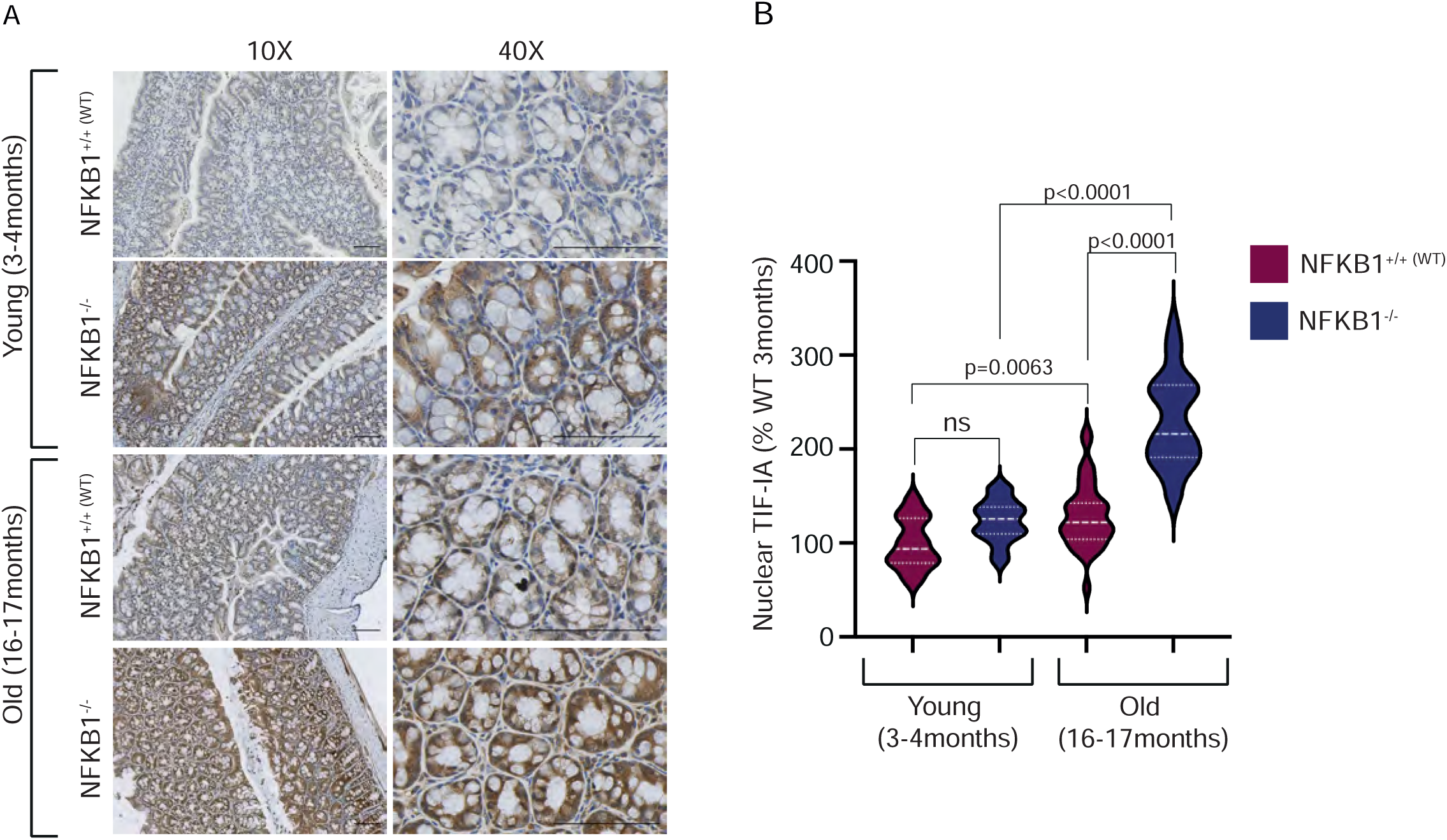
TIF-IA accumulates in the colon with age. Colonic sections from young and old wild type (WT) and NFKB1 null mice were stained for TIF-IA by immunohistochemistry (n=5 mice per group). Slides were scanned, five randomly selected regions of interest (ROI) across each colon identified, then the nuclear TIF-IA intensity of all cells within each ROI determined using QuPath digital pathology software. Sections shown were all stained simultaneously, and images (10X and 40X magnification) are representative. Scale bars=100uM. P values were obtained using a Tukey multiple comparison test, as described in the methods section.

## Discussion

Here we demonstrate that the PolI complex component, TIF-IA, acts as a key switch in the senescence regulatory network, inducing morphological changes to nucleoli, initiating the SASP and reinforcing the senescence phenotype. We demonstrate age-related accumulation of TIF-IA and identify p62 as a key regulator of TIF-IA stability downstream of DNA damage response.

### TIF-IA, nucleolar fusion and NF-κB pathway activation

Chronic inflammation is a hallmark of ageing and contributes to the pathophysiology of many age-related disorders such as diabetes, obesity, Alzheimer’s disease and cancer(51). Senescent cells, that accumulate in ageing tissue contribute to this chronic inflammatory state by secreting a cocktail of cytokines, chemokines and inflammatory factors (the SASP)(9,13). NF-κB is a key driver of the SASP and is also the transcription factor most closely associated with ageing(7,52). While progress has been made in understanding SASP reinforcement (13), the initial triggers of the NF-κB pathway in senescence are not fully understood. Another hallmark of ageing is enlarged nucleoli. In senescent cells, this enlargement is caused by nucleolar fusion. However, the impact of this fusion on ageing, inflammation and disease is also poorly characterised.

In this paper, we demonstrate that TIF-IA is required for both nucleolar fusion and the SASP in OIS and TIS. One possible explanation for our data is that the protein independently regulates these two hallmarks of senescence. However, in this and previous studies we have shown that in all contexts associated with nucleolar fusion (oncogene and DNA-damaged induced senescence, exposure to stress, ceramide, CDK4 inhibitor, hydrogen peroxide,) the NF-κB pathway is activated, as determined by degradation of IκB and/or transcription of NF-κB target genes(22,33). We have also shown that early changes in nucleolar morphology precede NF-κB pathway activation in senescence and stress response. Furthermore, we have previously demonstrated that in stress response, blocking TIF-IA degradation not only blocks nucleolar fusion but also NF-κB pathway activation(33). Similarly, here we show that blocking TIF-IA accumulation in senescence blocks both altered nucleolar morphology and the SASP, while us and others have demonstrated overexpressing TIF-IA causes nucleolar enlargement and transcription of SASP factors(24). Based on these data, we suggest the intriguing possibility that, rather than TIF-IA acting independently on nucleolar fusion and the SASP, TIF-IA-mediated nucleolar fusion lies upstream of NF-κB pathway activation and plays a role in initiating the SASP.

The mechanisms by which TIF-IA causes nucleolar fusion, and how this may lead to NF-κB pathway activation, have not yet been explored. Dogma would suggest that nucleolar size is directly linked to the rate of ribosome biogenesis. Indeed, there are many reports suggesting increased nucleolar area following senescence induction is caused by increased rDNA transcription and/or altered processing(18,31,43,44,53). For example, Nishimura et al used TIF-IA overexpression as a tool to accelerate ribosome biogenesis in mouse embryo fibroblasts (24). Similar to our findings, they demonstrated that TIF-IA overexpression caused increased nucleolar area, induction of SASP factors and senescence. They concluded that TIF-IA effects on rDNA transcription led to cell cycle arrest and senescence (they did not address links to the SASP). Here we found that following senescence induction, nuclear/nucleolar accumulation of endogenous TIF-IA has context dependent effects on rDNA transcription. That is, in some cell types/models of senescence, TIF-IA accumulation caused an increase in 47S transcription while in others, it was associated with a decrease. However, in all models we tested, TIF-IA accumulation was causally linked to changes to nucleolar morphology, the SASP and senescence. Similarly, in response to stress we have demonstrated that nucleolar fusion is associated with reduced rDNA transcription and NF-κB pathway activation(33). Therefore, we conclude that the mechanism by which TIF-IA controls nucleolar fusion and the SASP is independent of its role in PolI-driven transcription. Caragine et al. recently made the interesting observation that the nucleoplasm has an active (ATP dependent) contribution to nucleolar fusion and that disturbing chromatin dynamics altered nucleolar number and size(54). Depletion of components of the nuclear membrane LINC (linker of nucleoskeleton and cytoskeleton) complex and damage of rDNA have also been shown to regulate nucleolar number and morphology(55–57). Future studies will involve exploration of the role of TIF-IA in these alternative pathways to nucleolar fusion. Our findings that degradation and accumulation of TIF-IA have similar effects on nucleoli and NF-κB would suggest that, whatever the mechanism, there is a sweet spot of TIF-IA protein that prevents activation of this pathway.

One possibility to explain the link between nucleolar fusion and NF-κB pathway activation is nuclear displacement of the DNA sensor, cyclic GMP-AMP synthase (cGAS). The cGAS-Sting signalling pathway has emerged as a key regulator of NF-κB activity and the SASP (14,15). Although cGAS was originally identified as a cytoplasmic DNA sensor, it is now recognised that the majority of the protein resides in the nucleus, tightly bound to nucleosomes in an inactive state(58,59). Paradigm indicates that during senescence, genotoxic stress and breakdown of the nuclear membrane allow chromatin fragments to leak into the cytoplasm where they activate cGAS to initiate the SASP(13,60) (15). Given the gross chromatin changes associated with senescence and nucleolar fusion(54,61), and the fact that TIF-IA accumulates in the nucleoplasm as an early response to senescence, we suggest that TIF-IA either directly, or indirectly through nucleolar fusion/nuclear membrane breakdown, allows cGAS activation. Another nuclear protein involved in activation of NF-κB in response to DNA damage is IKKγ {Tufan, 2022 #1587} (62,63). It is now of paramount importance to define the mechanisms that links TIF-IA, nucleoli and NF-κB so that means can be identified to intervene in this pathway to prevent the SASP, chronic inflammation and disease.

### P62 and TIF-IA stability

Data presented here on senescence and aging, and elsewhere on apoptosis, indicate that control of TIF-IA protein stability is crucial for maintaining cellular homeostasis(33,64,65). Us and others have demonstrated that TIF-IA is degraded via the proteasome and it has been suggested that MDM2 acts as an E3 ligase in this process, ubiquitinating the N-terminal 94 amino acids of the protein(66). Here we make the novel observation that in steady state, TIF-IA is targeted for autophagosomal degradation by the cargo receptor, p62. Both the PB1 and ubiquitin binding domain of p62 are required for this interaction, suggesting that p62 binds to a ubiquitinated form of the protein. However, the N-terminal 94 amino acids of TIF-IA are not required. Indeed, we found p62 binds to TIF-IA 94-204, suggesting that the ubiquitination site(s) critical for p62 binding is within this minimal domain. We also found that P62 was unable to bind to an S44 phospho-mimetic mutant of TIF-IA. Based on these data, we propose that phosphorylation of S44 blocks ubiquitination within the 94-204 domain, P62 binding and autophagic degradation.

Our data indicate that activation of the DNA damage response kinase, ATM, is associated with reduced TIF-IA-p62 binding-TIF-IA accumulation, while ATM inhibition is associated with increased TIF-IA-P62 binding-TIF-IA loss. Mayer et al have previously demonstrated that S44 of TIF-IA is targeted for phosphorylation by CDK2/cyclinE, downstream of mTOR activation, and dephosphorylated by PP2A(49). ATM is known to target both mTOR and CDK2/cyclinE(67,68). Therefore, we considered that that ATM activation may cause hyperphosphorylation of S44 and loss of p62 binding through the MTOR pathway. However, we have previously demonstrated that rapamycin, that inhibits mTOR, has no effect on TIF-IA protein levels. In contrast, inhibiting PP2A, which is negatively regulated by ATM(69), caused TIF-IA accumulation. Based on these data, we suggest that the mechanism by which ATM alters TIF-IA protein levels is through inhibition of PP2A. However, it is also possible that ATM modulates the TIF-IA ubiquitination machinery. Further understanding of the dynamic flux of TIF-IA between autophagosomal and proteasomal degradation, and the upstream signalling pathways that control ubiquitination/p62 binding, will reveal insight into the regulation of cellular homeostasis/aging and is the focus of future research.

### TIF-IA in ageing and disease

Colorectal cancer is one of the leading causes of cancer morbidity and mortality worldwide and advanced age is one of the most significant risk factors(70). Although the mechanisms by which age contributes to colon cancer remain unclear, accumulation of senescent cells and secretion of SASP factors has been implicated(71). Here we used chronologically aged mice and a genetic model of aging to show that TIF-IA levels increase in the colon with age.

Similar to our findings with senescence, we find this accumulation is particularly evident in the nucleus of both stromal and epithelial cells. Senescent cells are rarely observed in tissue in vivo and are difficult to detect(9). One study that quantified β-galactosidase staining in small intestinal tissue from elderly mice found they comprised around 3.3% of cells(72).

Here we see TIF-IA levels are raised with age in the majority of colonic epithelial cells. One explanation for this discrepancy is that the DNA damage-induced increase in TIF-IA does not always lead to full senescence, as measured by β-galactosidase activity but that it sensitises to senescence induction. Multiplex immunohistochemistry will help us determine the dynamic relationship between DNA damage, nuclear accumulation of TIF-IA, nucleolar fusion and secretion of SASP factors in vivo.

## Acknowledgements

We would like to thank Terje Johansen (UIT, Artic University of Norway) for kindly providing the GFP-p62 plasmid. Simon Wilkinson, Mihaela Bozic and Priya Hari (ECRC, University of Edinburgh) for providing tools, reagents, and advice, and C. Nicol for providing help with figure preparation.

