## Supplemental files for "Identification of a P62-TIF-IA axis that drives nucleolar fusion and the senescence associated secretory phenotype"

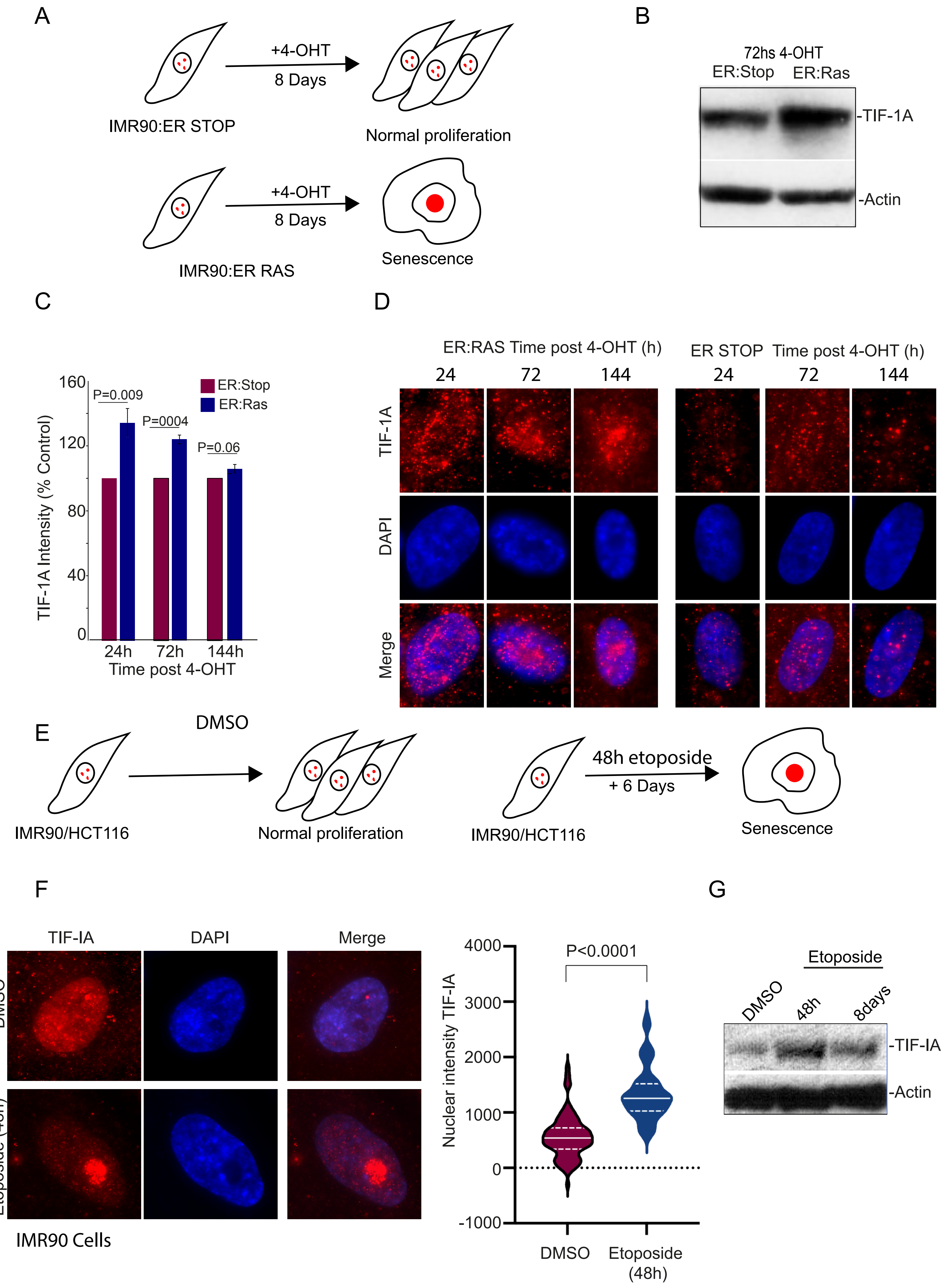

A
+4-OHT
8 Days
IMR90:ER STOP
Normal proliferation
+4-OHT
8 Days
IMR90:ER RAS
Senescence
B
72hs 4-OHT
ER:Stop ER:Ras
-TIF-1A
-Actin
C
160
120
80
40
0
TIF-1A Intensity (% Control)
P=0.009
P=0004
P=0.06
ER:Stop
ER:Ras
24h 72h 144h
Time post 4-OHT
D
ER:RAS Time post 4-OHT (h)
ER STOP Time post 4-OHT (h)
24 72 144 24 72 144
TIF-1A
DAPI
Merge
E
DMSO
IMR90/HCT116
Normal proliferation
48h etoposide
+ 6 Days
IMR90/HCT116
Senescence
F
TIF-IA
DAPI
Merge
DMSO
Etoposide (48h)
IMR90 Cells
4000
3000
2000
1000
0
-1000
Nuclear intensity TIF-IA
P<0.0001
DMSO
Etoposide (48h)
G
Etoposide
DMSO
48h
8days
-TIF-IA
-Actin

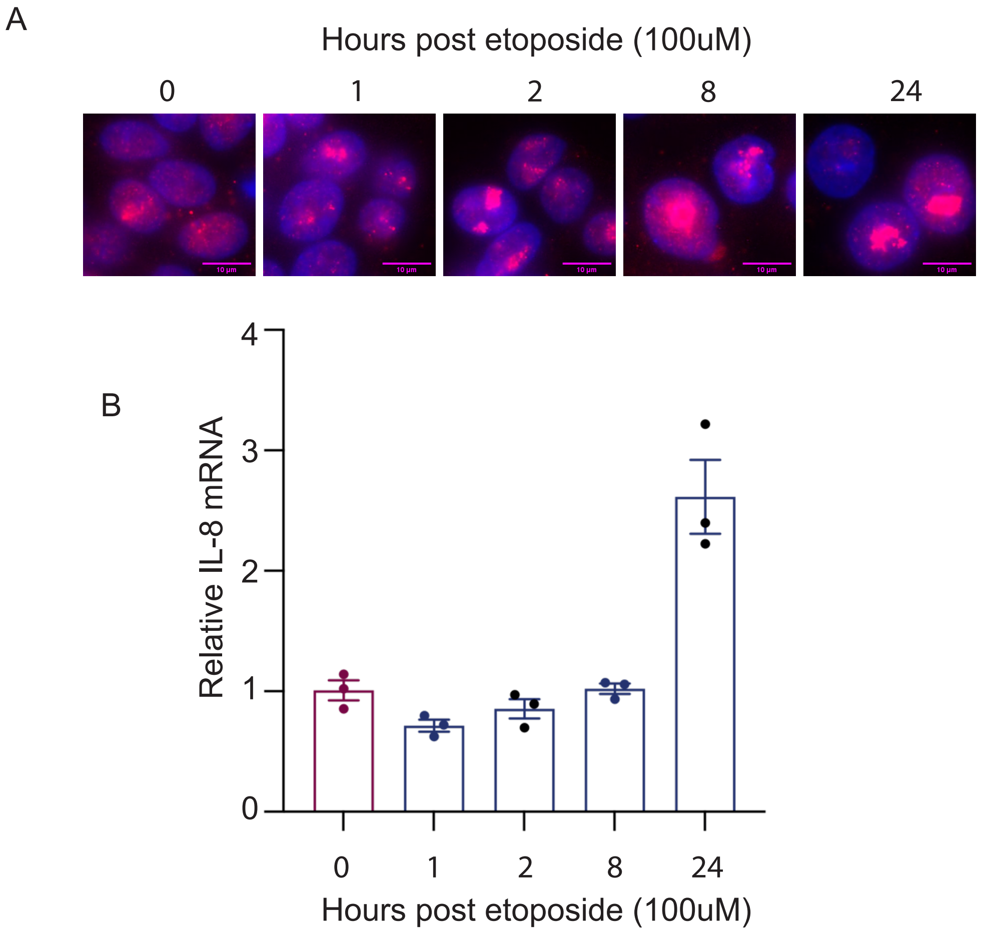

A
Hours post etoposide (100uM)
0
1
2
8
24
B
Relative IL-8 mRNA
4
3
2
1
0
0
1
2
8
24
Hours post etoposide (100uM)

**Supplemental figure legends**

**Supplemental Figure 1.** (A) Schematic outlining the model of oncogene-induced senescence (OIS) used. ER:RAS cells treated with 4-hydroxytamoxifen (4-OHT) undergo senescence while ER:STOP cells continue to proliferate. (B) Immunoblot showing total cell levels of TIF-IA in ER:STOP and ER:RAS cells treated for 72hours with 4OHT. (C and D). IMR90 ER:STOP or ER:RAS cells were treated with 4OHT in time course studies. (C) Anti-TIF-IA immunoblots were performed on whole cell lysates. FIJJI was used to quantify TIF-IA band intensity relative to actin (see Fig 1B for a representative image). N=3 (D) Representative immunomicrohraphs. Image quantification can be found in Fig. 1C. N=5. (E) Schematic outlining the model of therapy-induced senescence (TIS) used. (F and G) IMR90 cells were treated with DMSO or etoposide for the time specified. (F) Immunomicrographs demonstrating the localisation and intensity of TIF-IA. FIJJI was used to measure nuclear TIF-IA intensity. Five fields of view per experiment (at least 150 cells) were captured. N=5. (G) immunoblot showing whole cell levels of TIF-IA.

**Supplemental Figure 2**. A and B. HCT116 cells were treated with DMSO (0h) or etoposide (100uM) for the times shown. N=3 (A) Immunomicrographs show the localisation and intensity of TIF-IA. (B) qRT-PCR for the NF-κB target gene and SASP factor, IL-8. P values derived using a student T test.

Supplemental table 1- TIF-IA interactome

| Gene symbol | logFC Con | p.mod Con | logFC Etop | p.mod Etop |
|---|---|---|---|---|
| RRN3 | 12.5910436 | 5.14E-09 | 13.0386446 | 9.78E-09 |
| GFPT1 | 9.74304885 | 2.21E-08 | 10.2728212 | 1.74E-07 |
| SQSTM1 | 8.7237287 | 1.24E-07 | 8.45375894 | 1.99E-07 |
| NUDT21 | 7.15820014 | 1.28E-07 | 7.42850867 | 6.11E-06 |
| ITCH | 8.46412738 | 2.34E-07 | 8.50636131 | 1.42E-07 |
| SHKBP1 | 6.82024928 | 2.36E-07 | 6.69049496 | 6.75E-06 |
| ACOX1 | 5.47434283 | 4.45E-07 | 5.64392264 | 2.74E-06 |
| DLAT | 5.91302839 | 5.23E-07 | 6.06348075 | 9.87E-06 |
| IGKV1-6 | 8.11709911 | 5.75E-07 | 8.41142833 | 1.05E-06 |
| CPSF6 | 7.85386779 | 6.04E-07 | 7.82749411 | 4.59E-06 |
| CPSF7 | 5.31061417 | 9.94E-07 | 4.78398773 | 3.46E-05 |
| CD44 | 7.92801544 | 1.48E-06 | 6.81132229 | 2.23E-05 |
| MRPS14 | 4.40635351 | 2.13E-06 | 4.37763327 | 7.14E-05 |
| SAPCD2 | 6.428672 | 3.57E-06 | 6.56771679 | 4.44E-06 |
| PRDX4 | 4.88499713 | 3.89E-06 | 3.66982987 | 1.85E-05 |
| MRPL53 | 5.01940368 | 4.39E-06 | 5.33508813 | 3.24E-05 |
| TMEM160 | 5.69761276 | 5.88E-06 | 5.30146044 | 1.75E-05 |
| MRPL38 | 5.72105006 | 6.80E-06 | 3.58543799 | 0.02495734 |
| GOLGA5 | 4.73696803 | 7.24E-06 | 3.47138735 | 0.00051534 |
| PRRC2C | 4.05829638 | 8.13E-06 | 3.75820974 | 4.53E-05 |
| MRPL12 | 3.47630943 | 8.58E-06 | 3.2102411 | 0.00037951 |
| ENTR1 | 4.29306413 | 1.31E-05 | 3.47506376 | 0.00020068 |
| ZSCAN29 | 5.856472 | 2.28E-05 | 6.05446305 | 3.73E-05 |
| SPDL1 | 7.33656694 | 2.29E-05 | 7.1807598 | 4.73E-07 |
| KEAP1 | 6.32686686 | 3.40E-05 | 6.59520526 | 1.91E-06 |
| USP22 | 5.74860489 | 4.94E-05 | 6.08304576 | 3.94E-05 |
| RACGAP1 | 3.35338792 | 5.01E-05 | 2.93392106 | 3.79E-05 |
| WWP2 | 6.66087812 | 5.96E-05 | 6.7528646 | 2.74E-06 |
| C4A | 5.71781215 | 0.000137 | 5.81351466 | 2.20E-05 |
| MRPL43 | 3.04503328 | 0.00032973 | 3.73093641 | 0.0005896 |
| RBM26 | 2.74542665 | 0.00039366 | 2.8163734 | 8.33E-05 |
| DBT | 1.6527997 | 0.00043165 | 1.60875041 | 0.01146809 |
| RPL15 | 1.35812557 | 0.00052364 | 1.16568044 | 0.00369707 |
| RPL14 | 1.10365372 | 0.00104547 | 1.50539983 | 0.0070358 |
| PKM | 1.46419943 | 0.00105955 | 1.48906473 | 0.00395261 |
| CARM1 | 4.99911388 | 0.00156681 | 6.04252211 | 5.48E-06 |
| KHDRBS1 | 1.7865935 | 0.00196634 | 1.70311115 | 0.00104554 |
| MRPS34 | 1.2786372 | 0.00201177 | 1.12870614 | 0.00385384 |
| EWSR1 | 4.2018257 | 0.00216592 | 3.76540165 | 0.00101713 |

| | | | | |
|---|---|---|---|---|
| RPL18A | 1.44894302 | 0.00226716 | 1.64610042 | 0.00075196 |
| RPL7 | 1.1796517 | 0.00318085 | 1.15253996 | 0.00603102 |
| FASN | 1.10733179 | 0.00390274 | 0.98863489 | 0.01968386 |
| MISP | 1.94146957 | 0.00391191 | 1.96683714 | 0.00084272 |
| KIF23 | 4.06605417 | 0.00492501 | 2.88754336 | 0.00149669 |
| CHAMP1 | 3.89241198 | 0.00621866 | 3.71178149 | 0.0013401 |
| AGAP3 | 3.32400531 | 0.00731162 | 4.86936674 | 1.62E-05 |
| RBM27 | 2.81528866 | 0.01065905 | 1.18718134 | 0.01174521 |
| MRPS22 | 4.00804673 | 0.01066076 | 1.7564696 | 0.00052819 |
| DHX33 | 2.85445495 | 0.01092211 | 3.76689929 | 0.00531592 |
| NIPSNAP2 | 3.8231264 | 0.01187171 | 5.74617978 | 0.00719359 |
| FNBP4 | 4.64949418 | 0.01229048 | 4.80408746 | 0.01711687 |
| KCTD3 | 5.31783379 | 0.01569304 | 6.9366733 | 2.90E-05 |
| MRPS26 | 3.74422455 | 0.01570617 | 3.69134592 | 0.00198682 |
| PDHB | 2.91365341 | 0.01610952 | 4.90453783 | 0.01866498 |
| EXOSC4 | 5.41593231 | 0.032756 | 5.14377424 | 0.0366667 |
| PIP | 3.51593865 | 0.03434565 | 3.45879411 | 0.05202691 |
| MRPS9 | 2.78109902 | 0.04410336 | 1.72002569 | 0.0013209 |
| PDHX | 4.09483108 | 0.04414668 | 3.67644819 | 0.00057216 |
| C3 | 6.24991503 | 3.15E-06 | N/A | N/A |
| TRIP6 | 3.58831444 | 1.69E-05 | N/A | N/A |
| PSMC6 | 4.33571731 | 3.23E-05 | N/A | N/A |
| YLPM1 | 3.08677479 | 4.40E-05 | N/A | N/A |
| MRPS18B | 3.38027285 | 0.00012024 | N/A | N/A |
| RPL6 | 1.26166845 | 0.00059029 | N/A | N/A |
| HSPA5 | 1.28475338 | 0.00221786 | N/A | N/A |
| DBN1 | 1.36398152 | 0.00634858 | N/A | N/A |
| PML | 2.13123727 | 0.00773501 | N/A | N/A |
| ZBTB2 | 3.82845719 | 0.02967962 | N/A | N/A |
| PRMT5 | 1.93539649 | 0.03349005 | N/A | N/A |
| HNRNPH2 | 1.52957287 | 0.03962106 | N/A | N/A |
| KRT2 | 1.55152765 | 0.04401369 | N/A | N/A |
| DSTN | N/A | N/A | 4.81186434 | 0.00012127 |
| EXOSC2 | N/A | N/A | 5.27719971 | 9.57E-06 |
| GNL2 | N/A | N/A | 3.85279081 | 0.04626468 |
| KRT77 | N/A | N/A | 4.1896462 | 0.04818267 |
| MRPS31 | N/A | N/A | 1.41568711 | 0.00150735 |
| NDUFS6 | N/A | N/A | 4.36409248 | 0.00033826 |
| NIPSNAP1 | N/A | N/A | 5.95724413 | 0.03491404 |
| PDHA1 | N/A | N/A | 1.82835433 | 0.00838029 |
| RPL10 | N/A | N/A | 1.10300522 | 0.0097007 |
| RPL4 | N/A | N/A | 1.05382545 | 0.00854217 |
| RPRD2 | N/A | N/A | 4.09767203 | 1.49E-05 |

| | | | | |
|---|---|---|---|---|
| RPS26 | N/A | N/A | 1.17195139 | 0.01561151 |
| RPS27A | N/A | N/A | 1.75844527 | 0.003589 |
| RPS8 | N/A | N/A | 1.38741395 | 0.01499743 |
| SKI | N/A | N/A | 4.37506628 | 0.01437793 |
| TPI1 | N/A | N/A | 1.28223981 | 0.0516085 |
| VASP | N/A | N/A | 2.98773654 | 0.01038667 |
| WTAP | N/A | N/A | 3.67933 | 0.02998733 |
| YTHDC1 | N/A | N/A | 3.35851261 | 0.05129819 |

***Supplemental Table 1***

HCT116 cells were treated with DMSO (con) or 100uM etoposide (Etop) for 8h. TIF-IA was immunoprecipitated from whole cell lysates using IgG or a specific TIF-IA antibody. Quantitative M/S  was then used to identify proteins that specifically interact with TIF-IA for each treatment condition (IgG v TIF-IA LogFC>2 pMod<0.05).
